# Lysosome-acidifying nanoparticles rescue A30P α-synuclein induced neuronal death in cellular and *Drosophila* models of Parkinson’s disease

**DOI:** 10.1101/2024.04.19.590288

**Authors:** Chih Hung Lo, Mengda Ren, Gavin Wen Zhao Loi, Eka Norfaishanty Saipuljumri, Jonathan Indajang, Kah Leong Lim, Jialiu Zeng

## Abstract

Parkinson’s disease (PD) is an age-related neurodegenerative disease characterized by histopathological hallmarks of Lewy bodies formed by accumulation of α-synuclein (αSyn) and progressive loss of dopaminergic neurons in the substantia nigra *pars compacta* of the midbrain, with clinical symptoms of motor deficits. Toxic protein accumulation of αSyn in PD is associated with autolysosomal acidification dysfunction that contributes to defective autophagy-lysosomal degradation system. While lysosome-acidifying nanoparticles have been applied as therapeutics to ameliorate dopaminergic neurodegeneration in neurotoxin mediated or αSyn aggregates induced mouse model of sporadic PD, lysosome-targeted approach has not yet been applied in synucleinopathy models of familial PD. Here, we report the first application of the new poly(ethylene tetrafluorosuccinate-co-succinate) (PEFSU)-based acidic nanoparticles (AcNPs) in A30P αSyn overexpressing SH-SY5Y cells and *Drosophila* models of PD. In the cellular model, we showed that AcNPs restore lysosomal acidification, promote autophagic clearance of αSyn, improve mitochondrial turnover and function, and rescue A30P αSyn induced death in SH-SY5Y cells. In the *Drosophila* model, we demonstrated that AcNPs enhance clearance of αSyn and rescue dopaminergic neuronal loss in fly brains and improve their locomotor activity. Our results highlight AcNPs as a new class of lysosome-acidifying therapeutic for treatment of PD and other proteinopathies in general.

## Introduction

Parkinson’s disease (PD) is the second most common neurodegenerative disorder in the world, afflicting more than 10 million people (1, 2). Histopathological hallmarks of PD include the accumulation of α-synuclein (αSyn) aggregates in Lewy bodies within vulnerable neurons and the progressive loss of dopaminergic neurons in the substantia nigra *pars compacta* of the midbrain (3, 4). Clinical symptoms of PD include motor deficits such as tremor, rigidity, bradykinesia, and postural instability (5). The majority of PD cases are sporadic, with approximately 10-15% being familial (6). Both missense mutations and increased copy number such as duplication or triplication of the *SNCA* gene encoding αSyn cause early onset autosomal dominant PD (7). Even in patients without history of genetic inheritance, multiple genome-wide association studies have established that variations at the *SNCA* locus contribute significantly to the etiology of sporadic disease (7, 8), further suggesting the significance of targeting αSyn variants associated toxic protein aggregation and accumulation.

Growing evidence has shown that autophagic and lysosomal dysfunction and associated impairments in the autolysosomal degradation pathway contribute to the pathogenesis of PD (9-13). For example, neuropathological analysis has shown the presence of αSyn accumulation in cellular organelles such as lysosomes in post-mortem PD brains (10-12). Moreover, the depletion of lysosomes in dopaminergic neurons results in reduced levels of lysosomal-associated proteins and an observed buildup of undegraded autophagosomes in post-mortem PD brain tissues and different mouse models of PD (11, 12). Autolysosomal dysfunction has also been observed in synucleinopathy-associated PD models such as A30P and A53T αSyn transgenic mouse models (14-16). *Drosophila* expressing human αSyn carrying the disease-linked mutations in a pan-neuronal pattern faithfully replicate human PD (17) and is an established model organism for studying the autolysosomal degradation system in neurodegenerative diseases (18). Specifically, in *Drosophila* overexpressing human A30P αSyn, the gene expression of a lysosomal H^+^-ATPase, which is responsible for maintaining lysosomal acidification, was repressed (19), suggesting defective lysosomal function. Studies have also shown that *Drosophila* overexpressing human wild-type (WT), A30P, or A53T αSyn develop adult-onset, progressive degeneration of dopaminergic neurons with widespread Lewy-body-like inclusions and show locomotor deficit as monitored by progressive loss of climbing ability (17). In αSyn induced cellular models of PD, αSyn aggregates accumulate in the lysosomes (20), induce lysosomal dysfunctions such as lysosomal acidification defect (21) and lysosomal membrane permeabilization (22), impair mitochondrial function (23-25), and lead to neuronal death (26).

Lysosomes play a key role in the regulation of cellular functions, governing both cell death and survival (27). As the final destination within the autolysosomal pathway, lysosomes are responsible for the completion of the cellular degradation processes. Maintaining an optimal pH level typically between 4.0 and 5.0 is crucial for enabling fusion with autophagosomes, thereby ensuring proper autolysosomal functions such as organelle formation, enzymatic activity, and effective clearance of cellular debris and toxic protein aggregates (27). While many therapeutic agents have been discovered to promote autophagic function (28-30), few has been developed to specifically target lysosomes. Lysosome-acidifying nanoparticles have emerged as tools to target and re-acidify impaired lysosomes with elevated pH. These nanoparticles localize into lysosomes and degrade to release their component acids, thereby lowering the lysosomal luminal pH. In the context of PD, poly(lactic-co-glycolic) acid-based nanoparticles with the capacity to acidify lysosomes have been used to restore lysosomal acidification in human patient-derived *ATP13A2* and *GBA* mutant fibroblasts (31), environmental toxin such as 1-methyl-4-phenyl-1,2,3,6-tetrahydropyridine (MPTP) induced cellular and mouse models of PD (31, 32), as well as mice injected with PD patient-derived Lewy body extracts carrying αSyn aggregates (33). However, there is a lack of studies showing whether re-acidifying of impaired lysosomes in genetic models of synucleinopathy-associated PD can restore autolysosomal function and rescue dopaminergic neuronal function and motor deficit.

In this study, we report the first application of a new type of lysosome-targeting poly(ethylene tetrafluorosuccinate-co-succinate) (PEFSU)-based acidic nanoparticles (AcNPs) to re-acidify impaired lysosomes and restore autolysosomal function in cellular and *Drosophila* models of PD with A30P αSyn overexpression. The synthesized AcNPs are spherical and around 100 nm in size as well as possess low polydispersity, high stability, and good localization into lysosomes. In A30P αSyn-overexpressing SH-SY5Y cells, lysosomal pH elevation and impaired autolysosomal function were evident, leading to αSyn accumulation, reduced mitochondrial turnover, and consequent cell death (**Fig. 1A**). We showed that AcNPs effectively re-acidify impaired lysosomes and restore autolysosomal function as well as enhance αSyn clearance and mitochondrial turnover and function, thereby attenuating A30P αSyn induced cell death (**Fig. 1B**). As a proof-of-concept *in vivo* study, we further demonstrated that AcNPs restore autolysosomal function and rescue dopaminergic neuronal death and locomotor deficit in *Drosophila* with human A30P αSyn overexpression. We propose targeting lysosomal acidification dysfunction as an effective therapeutic strategy for managing familial synucleinopathy related PD. Furthermore, we highlight AcNPs as a new class of lysosome-acidifying therapeutic with potential applications for treatment of PD and other proteinopathies.

**Fig. 1.**
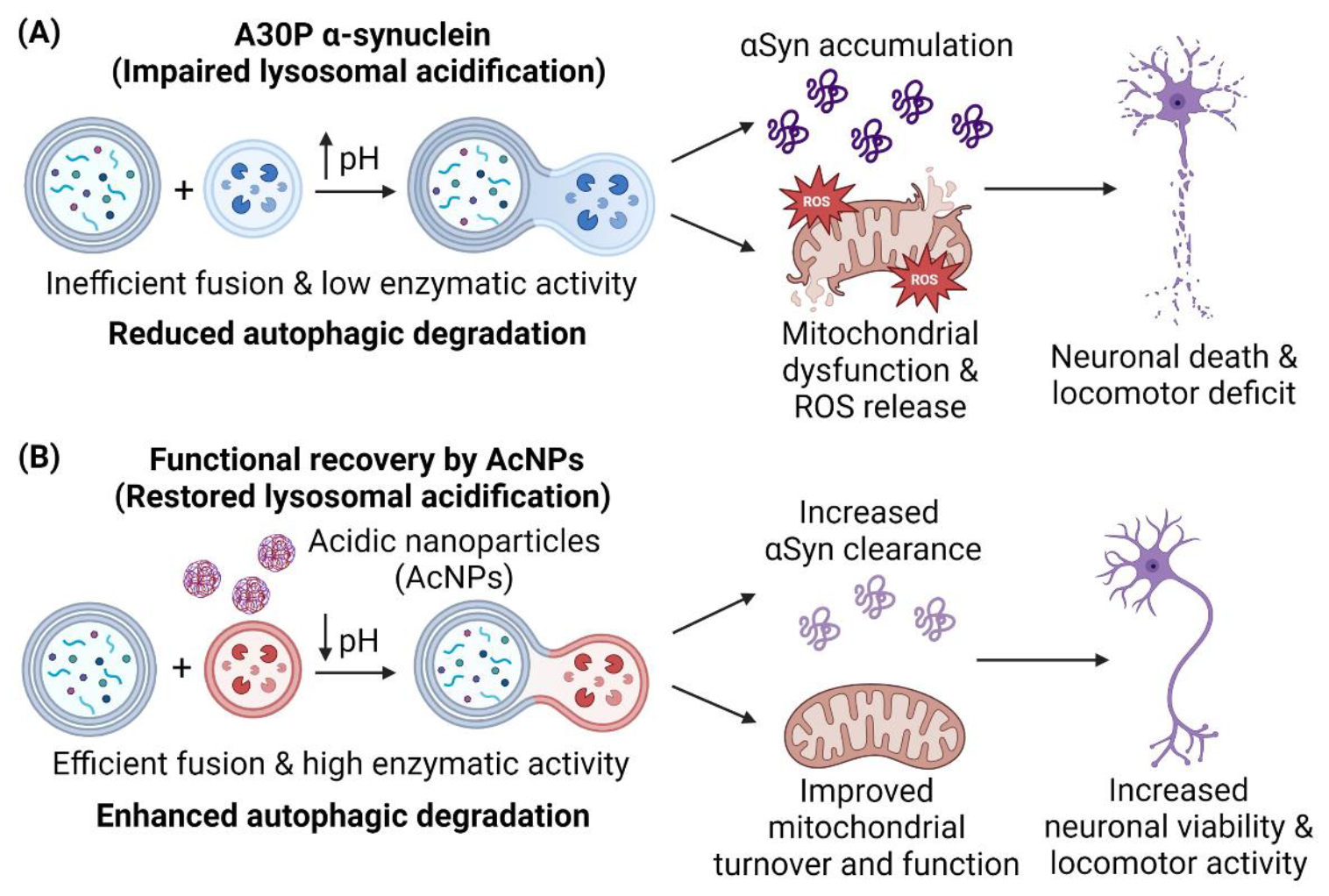
A30P αSyn induced neuronal death and locomotor deficit and functional recovery of neuronal functions by lysosome-targeting AcNPs. (A) Overexpression of A30P αSyn impairs lysosomal acidification, leading to inefficient fusion of autophagosomes and lysosomes, lowered lysosomal enzymatic activity, and overall reduced autophagic degradation. This results in αSyn accumulation as well as mitochondrial dysfunction and release of reactive oxygen species (ROS), implicating neuronal death and locomotor deficit. (B) Treatment of AcNPs re-acidify impaired lysosomes and promote their functions in neurons which leads to enhanced autophagic degradation and clearance of αSyn as well as improved mitochondrial turnover and function, resulting in increased neuronal viability and locomotor activity.

## Materials and Methods

### MAcNPs synthesis and characterizations

The AcNPs were synthesized from poly(ethylene tetrafluorosuccinate-co-succinate) (PEFSU) polyesters through a nanoprecipitation method based on previously established protocol (34). Briefly, PEFSU 25:75 polyesters composing of 25% tetrafluorosuccinic acid and 75% succinic acid were dissolved in acetonitrile (Sigma Aldrich, Cat # 271004) and filtered through a 0.2 μm syringe filter (Millipore, Cat# Z741696) to remove any particles. The polyester solution in acetonitrile was slowly added drop by drop into the vigorously stirred aqueous sodium dodecyl sulfate (SDS) (Sigma Aldrich, Cat# L3771) solution at 1700 rpm. The resulting emulsion was dialyzed against MilliQ water in SnakeSkin dialysis tubing (MWCO 10 kDa) (Thermofisher Scientific, Cat# 68100) for 24 h. NP size was measured by dynamic light scattering using the Brookhaven 90 plus NP size analyzer. For scanning electron microscopy characterization, the AcNPs were applied onto silicon wafers and allowed to air dry overnight. The wafers were then attached to aluminum stubs using copper tape and sputter coated with a 5 nm-thick Au/Pd layer. Image acquisition was conducted using a Supra 55VP field emission scanning electron microscope (ZEISS) with an accelerating voltage of 2 kV and a working distance of 5.5 cm.

### NPs degradation assays

For analysis of chemical components upon NPs degradation, the NPs solution was centrifuged at specific time points and the pellet were dried overnight in N_2_ atmosphere. The pellet was then dissolved in tetrahydrofuran and filtered through a 0.22 μm syringe filter. Gel permeation chromatography was performed to determine the molecular weights of the NP polymers using tetrahydrofuran as the eluent. To determine how different pH conditions may affect NPs degradation and the extent of acidification, NPs at 10 mg/mL concentration were diluted in phosphate saline buffer (PBS) adjusted to pH 6.0 or 7.4. The pH changes due to NPs degradation are measured using a pH meter at different time intervals.

### Cell culture and cytotoxicity assay

SH-SY5Y cells (ATCC, Cat# CRL-2266) were cultured using DMEM/F12 media (Thermofisher scientific, Cat# 11320033) supplemented with 1% penicillin-streptomycin (Thermofisher Scientific, Cat# 15140122) and 10% fetal bovine serum (Gibco FBS HI, Cat# A5256701) in a CO_2_ incubation at 37 °C. The effect of NPs on cell viability was determined using an MTS Assay Kit (Abcam, Cat# ab197010). SH-SY5Y cells transfected with vector-only control or A30P αSyn were cultured in a 96-well plate at 15000 cells/well for 24 h followed by treatment of respective doses of the AcNPs for 24 h. The cell viability was quantified relative to the vector-only control.

### Lysosomal localization, pH measurement, and cathepsin activity assays

For determination of NPs uptake into cells and their localization in sub-cellular compartments, SH-SY5Y cells were first incubated with rhodamine labeled AcNPs for 24 hours followed by the addition of LysoTracker blue dye (Thermofisher Scientific, Cat # L7525) according to manufacturer’s protocol for 30 min. The cells were then incubated with fresh media and imaged using confocal microscope (ZEISS) to acquire the fluorescent images at blue (excitation/emission 405/460 nm) and red (excitation/emission 590/610 nm) channels. For lysosomal pH measurements, SH-SY5Y cells transfected with vector-only control or A30P αSyn with and without AcNPs treatment were stained with LysoSensor™ Yellow/Blue DND-160 (Thermofisher scientific, Cat# L7545) at 1 μM concentration for 15 min. Image acquisition was performing using confocal microscopy with excitation at 360 nm and emission at 440 nm (blue channel) and 540 nm (yellow channel) (35). Lysosomal pH quantification was conducted based on the ratio between yellow and blue intensity and benchmarked against a standard curve with known pH values. The standard curve was generated by matching the ratio of yellow and blue fluorescence from LysoSensor™ to known pH using 2-(N-morpholino) ethanesulfonic acid buffer of varying pH (36). To perform cathepsin D and L activity assays, SH-SY5Y cells under respective conditions were stained with cathepsin D and L kits (Abcam, Cat# ab65302 and ab270774) following company protocols. The cells were then washed with PBS and before taking fluorescence measurements using a Synergy H1 model of hybrid multi-mode plate-reader with excitation/emission 328/460 nm for cathepsin D and 592/628 nm for cathepsin L. The fluorescence was normalized to cell count for each condition.

### Lysosomal membrane permeabilization assay

To perform lysosomal membrane permeabilization assay, SH-SY5Y cells transfected with vector-only control or A30P αSyn with and without AcNPs were fixed using a 4% paraformaldehyde (VWR, Cat# ALFAJ61899.AK) in PBS solution for 20 min and washed with PBS and 0.1% Triton X-100 (Sigma Aldrich, Cat# X100) in PBS. The fixed cells were then blocked with 5% normal goat serum (NGS) (Thermofisher Scientific, Cat# 31872) for 1 h at room temperature. To probe for lysosomal membrane permeabilization, the cells were stained with primary antibodies against galectin-3 (Proteintech, Cat# 14979-1-AP) overnight at 4 °C, followed by incubation with Goat anti-Rabbit 488 nm (Thermofisher Scientific, Cat# A-11008) fluorescent secondary antibody for 1 h at room temperature. The cells were then washed with PBS and stained with DAPI (Thermofisher Scientific, Cat# 9542). Image acquisition was performed using confocal microscopy with blue (excitation/emission 405/460 nm) and green (excitation/emission 490/520) channels.

### Characterization of mitochondrial turnover, morphology, and functions

To determine mitochondrial turnover under different conditions, SH-SY5Y cells were co-transfected with the mitophagy reporter mCherry-GFP-FIS1 (University of Dundee, Cat# DU55501) and vector-only control or A30P αSyn for 24 h, followed by AcNPs treatment for another 24 h. Before image acquisition, the LysoSensor™ Blue DND-167 dye (Thermofisher Scientific, Cat# L7533) was added to the cells for 15 min. Fluorescence images were captured using confocal microscopy with blue (excitation/emission 405/460 nm), green (excitation/emission 490/520 nm), and red (excitation/emission 590/610 nm) channels. To characterize mitochondrial morphology, SH-SY5Y cells under respective treatment conditions were stained with MitoTracker Deep Red (Thermofisher Scientific, Cat# M22426) following manufacturer’s protocol for 30 min. Fluorescence images were captured using confocal microscopy with excitation/emission wavelengths of 640/665 nm. The acquired confocal images of mitochondria were processed and analyzed using the Mitochondrial Network Analysis (MiNA) ImageJ macro (37). Quantifiable measurements of mitochondrial morphology, such as mitochondrial footprint (μm^2^) and network branch length (μm), were plotted. Consistent processing parameters used for analysis were applied across images from different treatment conditions. To measure mitochondrial membrane potential and superoxide release, SH-SY5Y cells under respective treatments were cultured in 96-well plate, followed by analysis using the TMRE-Mitochondrial Membrane Potential Assay Kit (Abcam, Cat# ab113852) and the Mitochondrial Superoxide Assay Kit (Abcam, Cat# ab219943) following manufacturer protocols. The measurements were obtained using a Synergy H1 plate-reader with excitation/emission wavelengths of 549/575 nm for TMRE and 510/610 nm for the superoxide signals.

### Drosophila stocks and locomotor assay

*Drosophila* were cultured on standard dextrose-yeast-cornmeal food at 25 °C with 12 h cycles of light and dark. The *Drosophila* stocks used are from the Bloomington Drosophila Stock Center: P{GMR57C10-GAL4}attP40 (BDSC, Cat# 90854), P{UAS-Hsap\SNCA.A30P}40.1 (BDSC, Cat# 8147), and *w*^*1118*^ (BDSC, Cat# 5905). Adult male flies were grouped in batches of 20–30 per vial for aging. During the aging period, the food was replenished every two days for the first two weeks, and then replaced daily starting from the third week. DAM2 *Drosophila* activity monitors (TriKinetics) were used to measure the effect of A30P αSyn and AcNPs on their daily locomotor activities. To prepare *Drosophila* food for DAM2 characterizations, the fly food was dried and trimmed to a height of 5 mm. Then, 10 μL of AcNPs solution at concentrations of either 3 mg/mL or 30 mg/mL was evenly applied to the surface. After ensuring uniform coverage of the AcNPs on the food under a stereoscope, a section of food with AcNPs was inserted into the DAM2 glass tube, with the AcNPs facing the flies. Adult flies were anesthetized on the day of eclosure and individually placed into the DAM2 glass tube, and the opening is sealed with a piece of cotton. For locomotor assay, the DAM2 glass tube containing the flies and food were loaded onto the DAM2 *Drosophila* activity monitors to record the number of times that the flies cross from one end of the tube to another end. The recording was set at a frequency of 10 seconds and lasted for a period of 15 days. The recorded locomotor data were analyzed and presented using ImageJ plugin ActogramJ (38).

### Immunofluorescence characterizations of Drosophila brains

*Drosophila* brains at 5 weeks old were first fixed in 4% paraformaldehyde in PBS with 0.1% Triton X-100 on ice for 0.5 h, followed by fixing at room temperature for an additional 0.5 h. After washing, the fly brains were stained with primary antibody against tyrosine hydroxylase (TH) (Pel Freeze, Cat# P40101-0) for 24 h at 4 °C, followed by incubation with Goat anti-Rabbit 488 nm (Thermofisher Scientific, Cat# A-11008) fluorescent secondary antibody for another 24 h at 4 °C. The fly brain tissues were then incubated with DAPI (Thermofisher Scientific, Cat# 9542) for 3 hours, briefly washed, and mounted with Prolong Gold (Thermo Fisher Scientific, Cat# P36930) on bridged slides. The tissues were positioned with the anterior side facing upward, and a single layer of protocerebral anterior medial (PAM) cluster was exposed for TH staining. The TH-positive neurons within a single PAM cluster were imaged using confocal microscopy for both DAPI (excitation/emission 405/460 nm) and TH (excitation/emission 490/520) channels, with 512x512 resolution and 0.34 μm slice interval to generate z-stacks.

### Quantification of TH-positive neurons in Drosophila PAM cluster

Labkit training and prediction of TH-positive region were conducted on a computer featuring an NVIDIA Quadro P6000. Twenty representative z-stacks comprising of a spectrum of TH signals and backgrounds were merged vertically into a unified stack using ImageJ. The stacks then underwent >100 cycles of “scribbling-predicting-correcting” to train the TH-positive pixel classifier against backgrounds in Labkit. The resulting classifier file was applied consistently for subsequent analysis of TH-positive neurons. To identify TH-positive regions, the classifier file was used to batch process a folder of z-stack TH images with minimization of noise and vacuoles on the Labkit-predicted mask images using customized Matlab (Mathwork) code “Correct_Labkit_TH.m” to generate new masks. The new masks were then processed to fill in any small gaps or holes present within the TH-positive regions to ensure that the mask is continuous, resulting in a high-quality and refined mask suitable for subsequent image analysis. The processed mask images were imported into Imaris 8.4 as a new channel alongside the raw images. TH-positive surfaces and DAPI spots were created using Imaris in the Labkit channel and DAPI channel, respectively. The number of TH-positive neurons in the PAM cluster of fly brains was counted as DAPI spots located within the TH-positive surface by running the surface distance transformation extension outside the TH-positive surfaces.

### Western blotting

For preparation of cell lysates, SH-SY5Y cells under respective conditions were lysed using RIPA buffer (Thermofisher Scientific, Cat# 89900) supplemented with protease and phosphatase inhibitors (Thermofisher Scientific, Cat# 78440) as the lysis solution. Fly brain tissues were homogenized using the same lysis solution as described above. All lysates were kept on ice for 15-30 min followed by centrifugation at 13,500 g at 4 °C for 10 min. The total protein concentrations for each sample of the cell and tissue lysates were determined using the Pierce™ BCA Protein Assay Kit (Thermofisher Scientific, Cat# 23225). 4X Laemmli Sample Buffer (Bio-Rad, Cat# 1610747) with 2-mercaptoethanol (Sigma-Aldrich, Cat# M6250) was then added and the samples were boiled at 95 °C for 5 min before loading and running in 4-15% Mini-PROTEAN TGX Precast Protein Gels (Bio-Rad, Cat# 4561083) with Precision Plus Protein Dual Color Standards (Bio-Rad, Cat# 1610374). The proteins were transferred to polyvinylidene fluoride membrane (Millipore, Cat# ISEQ00010) following by incubation with blocking buffer (Bio-Rad, Cat# 1706404). The membranes were then incubated with respective primary and secondary antibodies before image acquisition using the ChemiDoc MP imaging system (Bio-Rad). Densitometry analysis was conducted using ImageJ and protein expression levels were normalized to α-tubulin loading control.

The list of primary antibodies used include Human α-synuclein (Abcam, Cat# ab212184 and BD Biosciences, Cat# 610787), human p62 (Cell Signaling Technology, Cat# 5114), human LC3 (Cell Signaling Technology, Cat# 12741), human α-tubulin (ProteinTech, Cat# 11224-1-AP), *Drosophila* Ref2P (Abcam, Cat# ab178440) and *Drosophila* α-tubulin (DSHB, Cat# 12G10). The secondary antibodies used are Goat Anti-Rabbit IgG H&L (HRP) (Abcam, Cat# ab6721), Goat Anti-Mouse IgG H&L (HRP) (Abcam, Cat# ab205719), and Horse Anti-Mouse IgG H&L (HRP) (Cell Signaling Technology, Cat# 7076). All antibody dilutions used were following company recommendations.

### Statistical analysis

Statistical analyses were conducted using the GraphPad Prism 9 software. Statistical analysis was performed by using one-way ANOVA with post hoc Tukey’s test for multiple comparisons. Statistical significance was determined by *P*<0.05 and indicated by \**P*<0.05, \*\**P*<0.01, \*\*\**P*<0.001, \*\*\*\**P*<0.0001 or ns for non-significance.

## Results

### AcNPs restore lysosomal acidification in A30P αSyn overexpressing SH-SY5Y cells

We first synthesized AcNPs that are capable of degrading in mildly acidic conditions and releasing their component acids to restore lysosomal acidification. The AcNPs formed are spherical and around 100 nm in size with low polydispersity and high zeta potential which indicates high stability (**Fig. 2A**). AcNPs illustrated a strong buffer acidification strength at mildly acidic environment of pH 6.0 but not at neutral condition of pH 7.4 (**Fig. S1A-C**). We also showed that AcNPs are nontoxic to SH-SY5Y cells (**Fig. S2A**), making them suitable to be used in restoring lysosomal acidification in the cells. Using rhodamine labeled AcNPs, we confirmed the uptake of AcNPs into SH-SY5Y cells and their sub-cellular localization into lysosomes (**Fig. 2B**). We then determined the extent of lysosomal acidification impairment associated with A30P αSyn overexpression. Overexpression of A30P αSyn elevated lysosomal pH from 4.6 to 5.2 as compared to the vector-only control in SH-SY5Y cells (**Fig. 2C-D**). Treatment of AcNPs at 50 μg/mL and 100 μg/mL lowered lysosomal pH from 5.2 to 4.7 in a dose-dependent manner (**Fig. 2C-D**). As a control, AcNPs did not alter basal lysosomal pH in SH-SY5Y cells transfected with vector-only control (**Fig. S2B**). Overexpression of A30P αSyn also induced lysosomal membrane permeabilization in SH-SY5Y cells which was rescued by AcNPs treatment (**Fig. 2E-F**). Furthermore, there was a reduction in lysosomal enzymes cathepsin D and L activities in A30P αSyn overexpressing cells and the enzyme activities were restored with treatment of AcNPs (**Fig. 2G-H**).

**Fig. 2.**
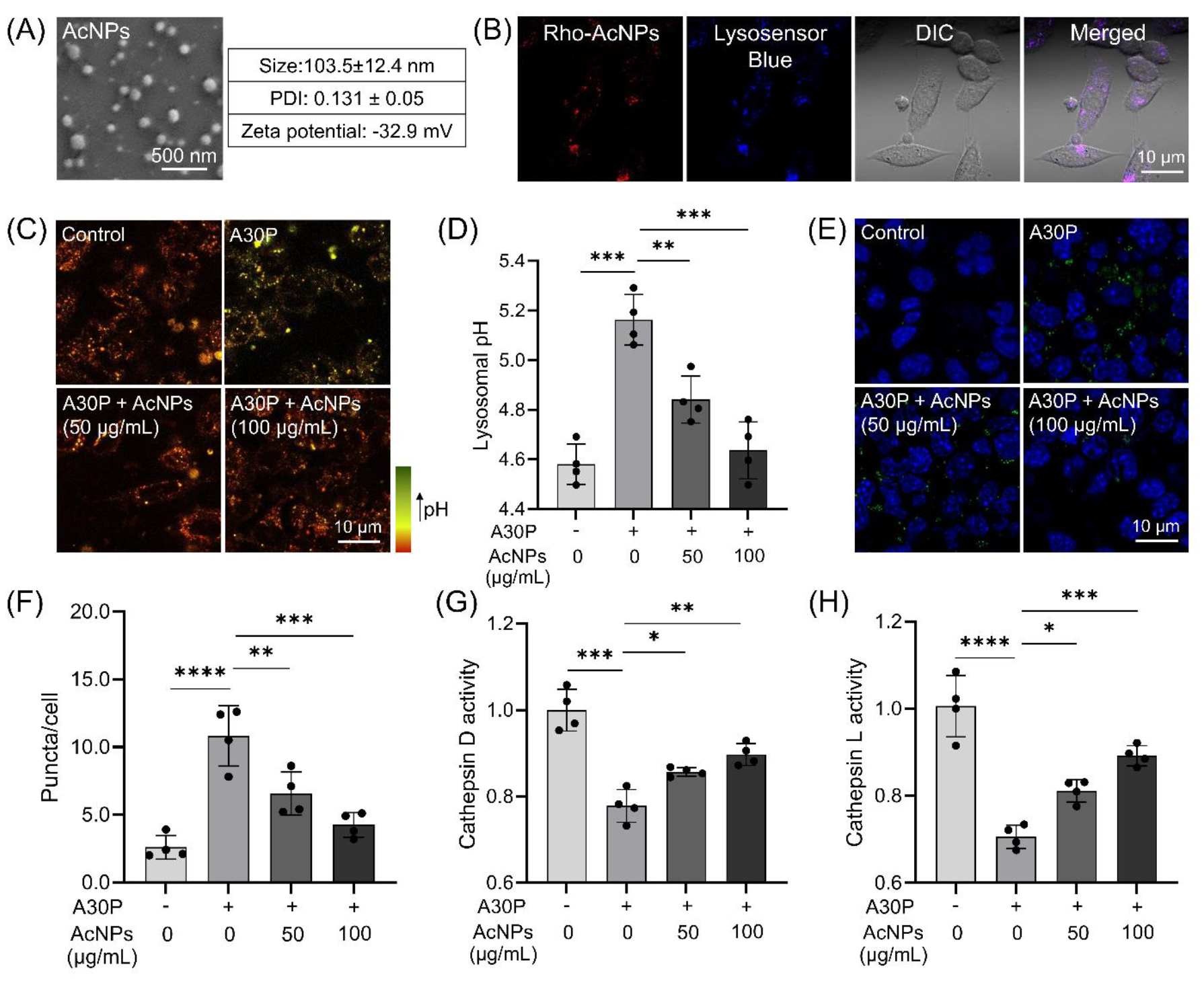
AcNPs restore lysosomal acidification and cathepsin activity in A30P αSyn overexpressing SH-SY5Y cells. (A) Characterization of lysosome-targeting AcNPs in terms of size, polydispersity index (PDI), and zeta potential. (B) Uptake of rhodamine labeled AcNPs into SH-SY5Y cells and their localization inside lysosomes. (C-D) Lysosomal pH measurement and quantification by confocal imaging with LysoSensor Yellow/Blue dye in A30P αSyn overexpressing SH-SY5Y cells with and without treatment of AcNPs. (E-F) Lysosomal membrane permeabilization as characterized by immunostaining with galectin-3 and image quantification in SH-SY5Y cells under respective treatment conditions. (G-H) Enzymatic activities of lysosomal cathepsin D and L in SH-SY5Y cells under respective treatment conditions. Data presented are relative to vector-only control. Data are means ± SD of N=4 independent experiments. \**P* < 0.05, \*\**P* < 0.01, \*\*\**P* < 0.001, and \*\*\*\**P* < 0.0001 using one-way ANOVA with post hoc Tukey’s test for multiple comparisons.

### AcNPs promote autophagic clearance of αSyn and improve mitochondrial turnover

Following the characterization of lysosomal acidification, we examined whether re-acidifying impaired lysosomes promotes autophagic degradation in A30P αSyn overexpressing SH-SY5Y cells. Treatment of AcNPs improves the autophagic degradation of autophagosome-associated proteins p62 and LC3II (39) as well as reduces αSyn accumulation (**Fig. 3A-D**). As functional autophagy is essential for the proper turnover of mitochondria, we then determined whether AcNPs improve mitochondrial turnover using a mCherry-GFP-FIS1 reporter co-expressed with A30P αSyn in SH-SY5Y cells. The mCherry-GFP-FIS1 reporter plasmid assesses the accumulation of mitochondria in lysosomes with varying levels of acidification by measuring fluorescence signals (40). The GFP fluorescence in the mCherry-GFP-FIS1 expressing cells will be quenched in sufficiently acidified lysosomes but not in poorly acidified lysosomes. The mCherry fluorescence remains stable as a control marker under all conditions to indicate the presence of mitochondrial protein. Overexpression of A30P αSyn led to an increase in the GFP fluorescence as compared to the vector-only control, indicating reduced lysosomal acidification (**Fig. 3E-F**). This results in the formation of white puncta arising from the colocalization of mitochondria (mCherry) in lysosomes (blue) with poor acidification (green). Treatment of AcNPs re-acidified impaired lysosomes and quenched GFP fluorescence, leading to the formation of purple puncta that arise from the colocalization of mitochondria (mCherry) in sufficiently acidified lysosomes (blue) (**Fig. 3E-F**). Overall, AcNPs promoted autophagic clearance of αSyn and improved mitochondrial turnover.

**Fig. 3.**
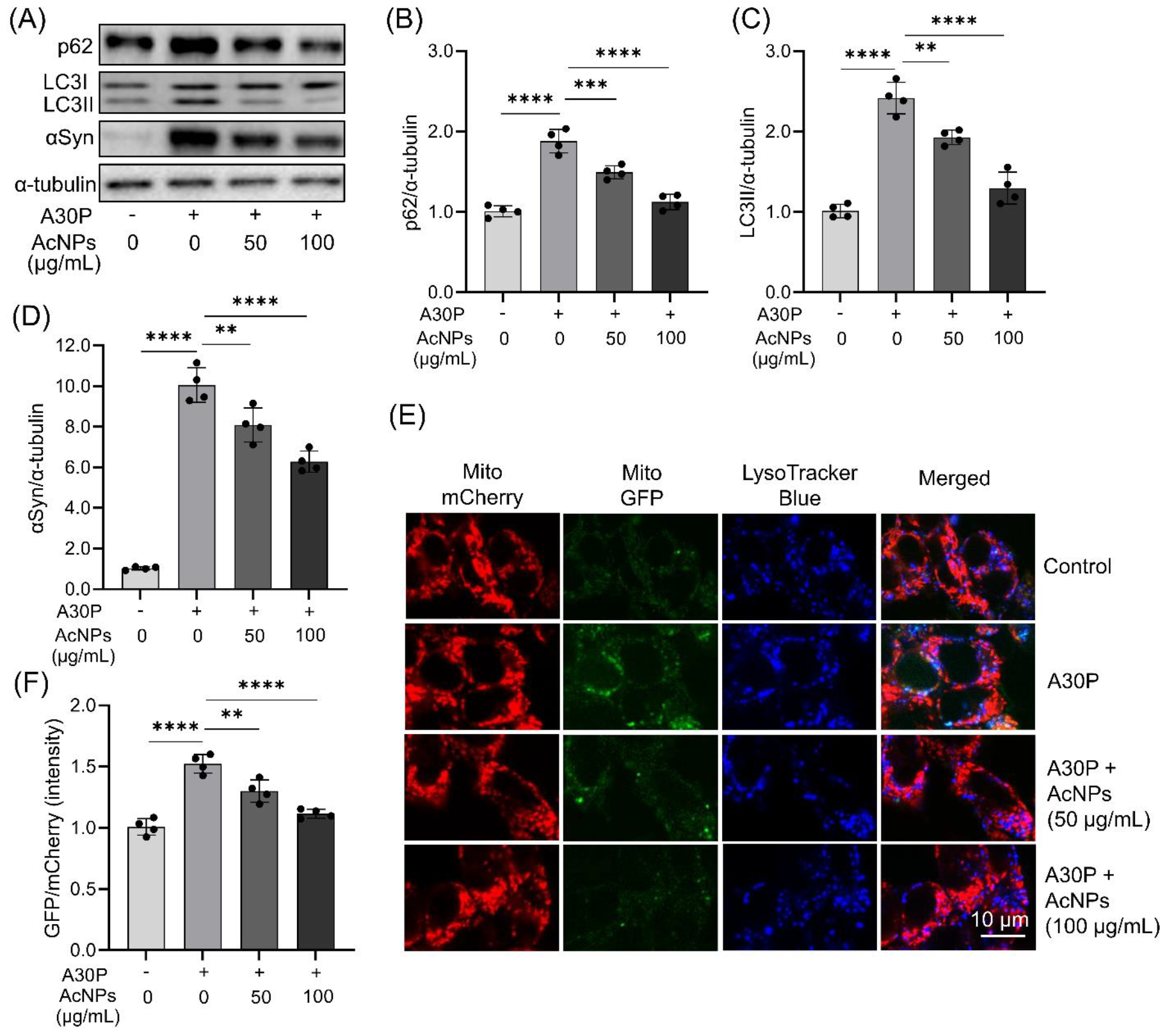
AcNPs promote autophagic degradation and improve mitochondrial turnover in A30P αSyn overexpressing SH-SY5Y cells. (A-D) Western blotting analysis and quantification of autophagy-associated proteins (B) p62 and (C) LC3II as well as (D) total αSyn in SH-SY5Y cells under respective treatment conditions with α-tubulin as a loading control. (E-F) Assessment of the extent of mitochondrial turnover under respective treatment conditions through confocal microscopy imaging and quantification of SH-SY5Y cells transfected with mCherry-GFP-FIS1 mitophagy reporter plasmid. Data presented are relative to vector-only control. Data are means ± SD of N=4 independent experiments. \*\**P* < 0.01, \*\*\**P* < 0.001, and \*\*\*\**P* < 0.0001 using one-way ANOVA with post hoc Tukey’s test for multiple comparisons.

### AcNPs restore mitochondrial functions and rescue A30P αSyn induced cell death

To further investigate whether improved mitochondrial turnover restores normal mitochondrial functions, we stained A30P αSyn overexpressing SH-SY5Y cells with MitoTracker Deep Red under AcNPs treated condition. First, overexpression of A30P αSyn increased mitochondrial fragmentation (**Fig. 4A-C**), which is indicative of mitochondrial impairment. Treatment of AcNPs increased the mitochondrial footprint and network branches, indicating that there is reduced mitochondrial fragmentation and increased function (**Fig. 4A-C**). Overexpression of A30P αSyn also decreased mitochondrial membrane potential (**Fig. 4D**) and elevated reactive oxygen species (**Fig. 4E**) which were reversed by AcNPs treatment. Furthermore, we demonstrated that AcNPs rescued A30P αSyn induced death in SH-SY5Y cells in a dose-dependent manner (**Fig. 4F**).

**Fig. 4.**
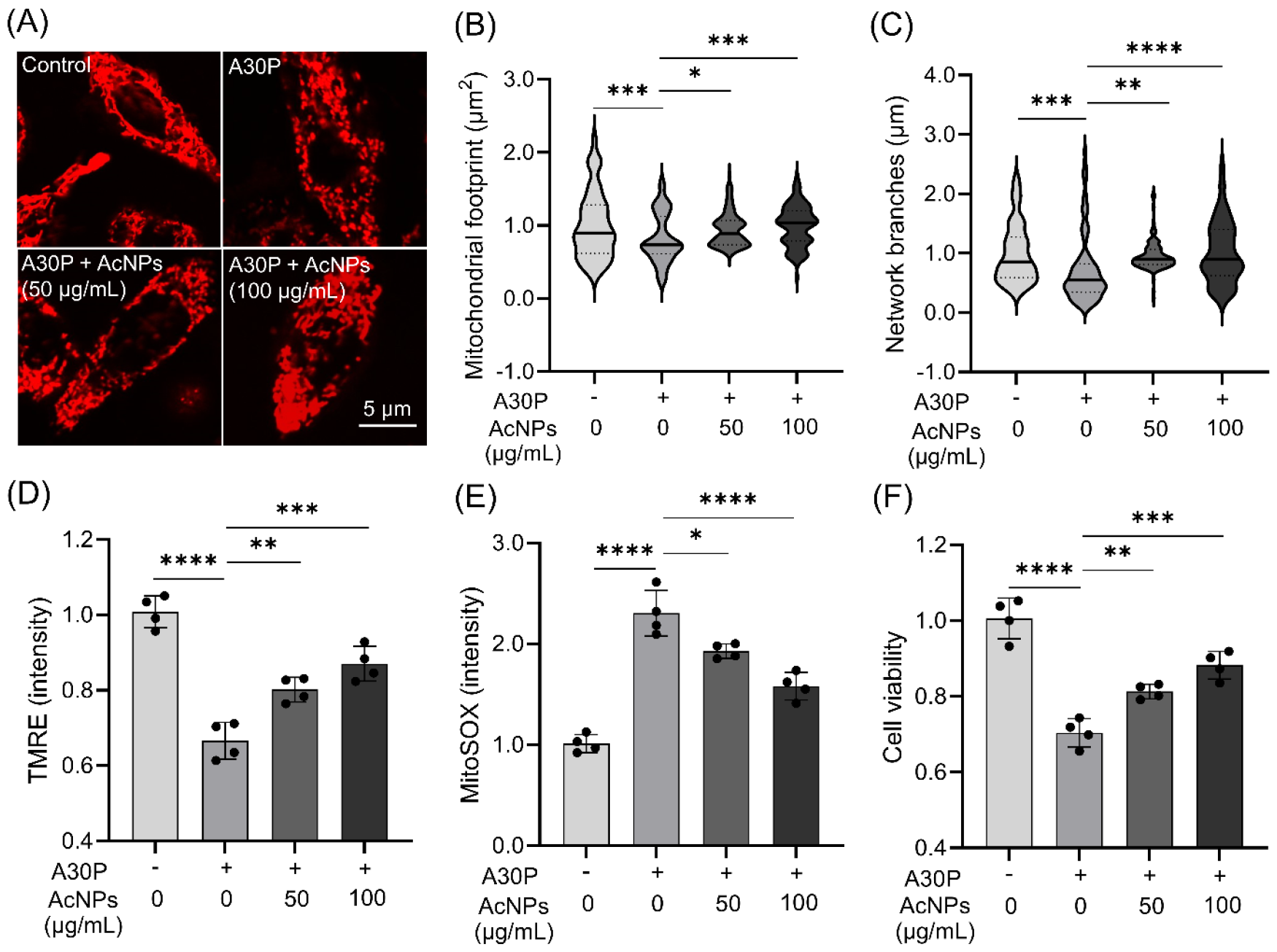
AcNPs enhance mitochondrial function and rescue A30P αSyn induced death in SH-SY5Y cells. (A) Assessment of mitochondrial morphology under respective treatment conditions through confocal microscopy imaging and quantification of SH-SY5Y cells stained with MitoTracker Deep Red. (B-C) Quantification of mitochondrial images using MiNA analysis to obtain measurements for (B) mitochondrial footprint and (C) network branches. (D-E) Assessment of mitochondrial functions through measurements of (D) mitochondrial membrane potential (TMRE) and (E) reactive oxygen species generation (MitoSOX) in SH-SY5Y cells under respective treatment conditions. (F) Measurement of cell viability of SH-SY5Y cells transfected with vector-only control or A30P αSyn and treated with and without AcNPs. Data presented are relative to vector-only control. Data are means ± SD of N=4 independent experiments. \**P* < 0.05, \*\**P* < 0.01, \*\*\**P* < 0.001, and \*\*\*\**P* < 0.0001 using one-way ANOVA with post hoc Tukey’s test for multiple comparisons.

### AcNPs promote αSyn clearance, rescue neuronal loss, and restore locomotor activity in A30P αSyn Drosophila

Having observed the effect of AcNPs in cells, we further investigated the effect of AcNPs in *Drosophila* with overexpression of human A30P αSyn in a pan-neuronal manner, which is a widely used model that has been shown to recapitulate key pathological features of human PD pathology (17, 18). First, to determine if there is impairment in the autophagic degradative function in *Drosophila*, we showed that there is an accumulation of autophagosome-associated protein Ref2P (fly orthologue of human p62), along with an elevated total αSyn level in the whole brain lysates of A30P αSyn *Drosophila* (**Fig. 5A-C**). Treatment with AcNPs reduced the accumulation of p62 and αSyn, indicating enhanced autophagic degradation and clearance of αSyn in fly brains (**Fig. 5A-C**). Next, we conducted immunofluorescence staining of fly brains using the TH marker to evaluate dopaminergic neuronal counts. A30P αSyn fly brains exhibited a significantly lower count of dopaminergic neurons compared to the control fly brains (**Fig. 5D-E**). AcNPs treatment rescued neuronal loss in A30P αSyn fly brains (**Fig. 5D-E**). Finally, overexpression of A30P αSyn reduced fly locomotor activity, which was restored by AcNPs treatment (**Fig. 6A-B**), demonstrating the *in vivo* efficacy of AcNPs.

**Fig. 5.**
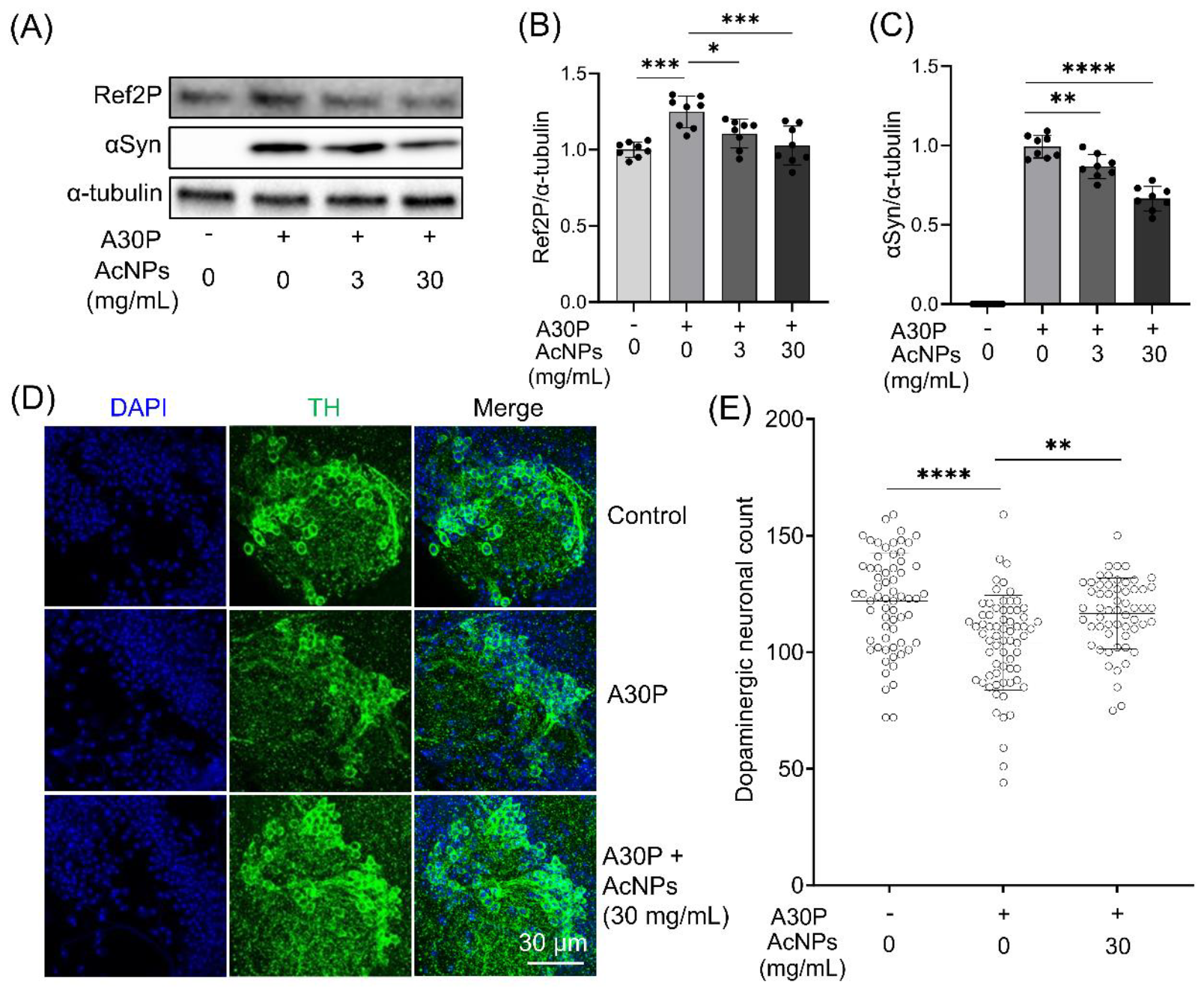
AcNPs promote αSyn clearance and rescue dopaminergic neuronal loss in A30P αSyn *Drosophila*. (A-C) Western blotting analysis and quantification of autophagy-associated proteins (B) Ref2P and (C) total αSyn in *Drosophila* overexpressing A30P αSyn under respective treatment conditions with α-tubulin as a loading control (N=8). (D-E) Immunofluorescence imaging and quantification of TH-positive dopaminergic neurons in the protocerebral anterior medial (PAM) of *Drosophila* brains under respective treatment conditions (N=57-67). \**P* < 0.05, \*\**P* < 0.01, \*\*\**P* < 0.001, \*\*\*\**P* < 0.0001, and ns indicates non-significance using one-way ANOVA with post hoc Tukey’s test for multiple comparisons.

**Fig. 6.**
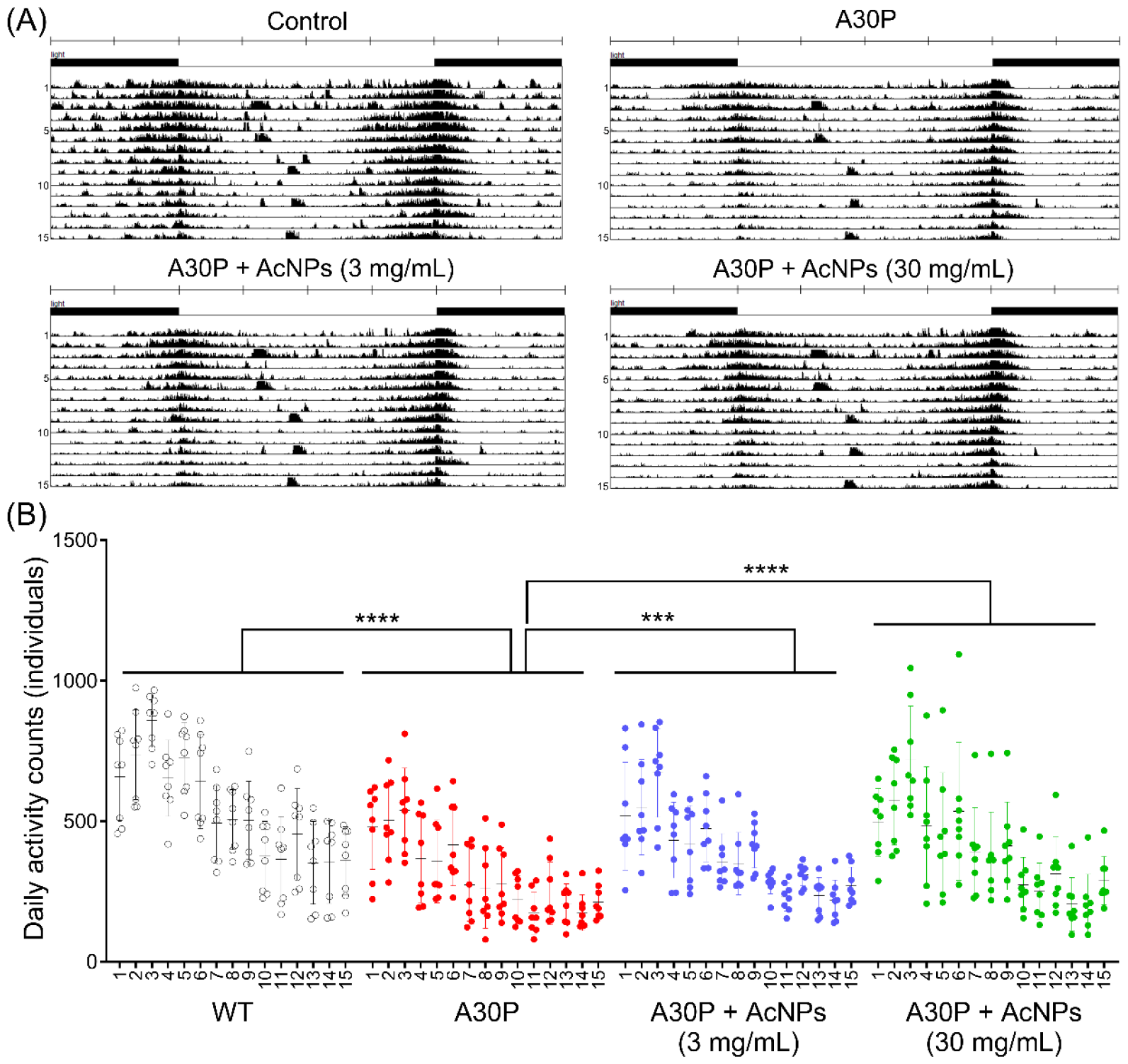
AcNPs restore locomotor activity in A30P αSyn *Drosophila*. (A) Mean actograms and (B) quantification of the daily locomotor activity counts of *Drosophila* under respective treatment conditions recorded using DAM2 *Drosophila* activity monitors (N=8). \*\*\**P* < 0.001 and \*\*\*\**P* < 0.0001 using one-way ANOVA with post hoc Tukey’s test for multiple comparisons.

## Discussion

In this study, we demonstrated for the first time the feasibility of targeting autolysosomal acidification dysfunction in A30P αSyn overexpressing cellular and *Drosophila* genetic models of PD. This is also a novel application of the new type of PEFSU-based lysosome-targeting AcNPs in PD models. We showed that AcNPs effectively re-acidified impaired lysosomes and enhanced autolysosomal activity in A30P αSyn-overexpressing SH-SY5Y cells, thereby promoting αSyn clearance, improving mitochondrial turnover and function, and rescuing A30P αSyn induced cell death. Importantly, we extended the application of AcNPs to *Drosophila* with human A30P αSyn overexpression to provide the first proof-of-concept *in vivo* rectification of autolysosomal dysfunction in a genetic model of synucleinopathy-associated PD. We demonstrated that AcNPs restore autolysosomal functions and attenuate dopaminergic neuronal loss in the fly brains, together with rescuing their locomotor deficits.

As *Drosophila* do not possess an orthologue of αSyn (18), flies ectopically expressing human WT or mutant αSyn provide established *in vivo* models for PD studies by recapitulating synucleinopathy related neuronal dysfunction and death as well as locomotor deficits (17). In *Drosophila*, several studies point to autophagic and lysosomal dysfunction in the presence of human WT or mutant αSyn overexpression, including the abnormal accumulation of autophagosomes (41-43) and defective mitochondrial turnover and function (19, 44). Specifically, our results are corroborated by a previous study using *Drosophila* overexpressing human A30P αSyn which illustrated the downregulation of genes responsible for lysosomal H^+^-ATPase and mitochondrial respiratory chain components, indicating the presence of autolysosomal acidification dysfunction and mitochondrial impairments (19). In terms of functional recovery, overexpression of LAMP2A (lysosomal protein) (43) and *Drosophila* homologue of DOR (autophagy regulator) (44) have been shown to increase autophagic activity and mitochondrial turnover as well as reverse loss of dopaminergic neurons and locomotor declines induced by the overexpression of human A30P αSyn in *Drosophila*. Our results complement these studies by highlighting the efficacy of AcNPs, as opposed to the overexpression of autolysosomal proteins, in facilitating functional recovery in A30P αSyn *Drosophila*.

Apart from lysosome-acidifying nanoparticles (27, 34, 36, 45, 46), there are several other strategies such as small molecules that target lysosomal proton pump and ion channels that have been developed to restore lysosomal acidification. For example, C381 activates lysosomal vacuolar H^+^-ATPase and promotes lysosomal acidification in neurotoxin MPTP treated mouse model of PD (47). ML-SA1 is an agonist of lysosomal channel transient receptor potential mucolipin 1 that has been shown to enhance lysosomal acidification and promote clearance of A53T αSyn in HEK293 cells and WT αSyn in immortalized human dopaminergic neurons (48). Other small molecules such as curcumin analog C1 and PF11 (27), which activate transcription factor EB to promote lysosome biogenesis and luminal acidification, have been applied to neurotoxins MPTP (49) and 6-hydroxydopamine (50) treated cellular and rat models of PD, respectively. Optimization of available lead compounds and nanomedicine as well as launching of new drug screen studies such as using artificial lysosomes loaded with trypsin (51, 52) may provide more effective lysosome-targeting therapeutics that can potentially be applied to PD. Furthermore, future studies may include direct *in vivo* monitoring of autolysosomal acidification dysfunction in PD using autolysosomal reporter flies (53) and mice (54).

The significance of targeting defective autolysosomal acidification is reflected by the notion of early impairment in autolysosomal acidification preceding neurodegeneration (27, 55-60). While additional evidence is necessary to strengthen the aspect of early impairment, there appears to be a positive correlation between disease penetrance and the degree of lysosomal acidification dysfunction in PD manifestations. For example, cells with A53T αSyn overexpression, which is a highly penetrant mutation (61, 62), show a more significant elevation of lysosomal pH compared to the less penetrant A30P or WT αSyn overexpression (21). It is also important to note that autolysosomal dysfunction has been associated with increased αSyn spreading in both genetic and sporadic models of PD (63, 64), thus contributing to disease progression. Taken together, targeting autolysosomal acidification dysfunction holds promise as an effective therapeutic strategy for treatment of PD and other proteinopathies in general.

## Authors’ contributions

J.Z. and C.H.L. conceived and directed the research project. J.Z. and C.H.L. conducted all cell-based experiments and performed data analysis. M.R. and L.K.L. carried out the *Drosophila* experiments and analyzed the data. G.W.Z.L. and E.N.S. provided technical support in cell culture and biochemical assays. J.I. assisted in image and data analysis. L.K.L. provided technical discussions and critical comments to the manuscript. J.Z. and C.H.L. wrote the manuscript. All authors approved the final version of the manuscript.

## Acknowledgements

The authors thank the funding sources for supporting this work.

---

**Fig. S1.**
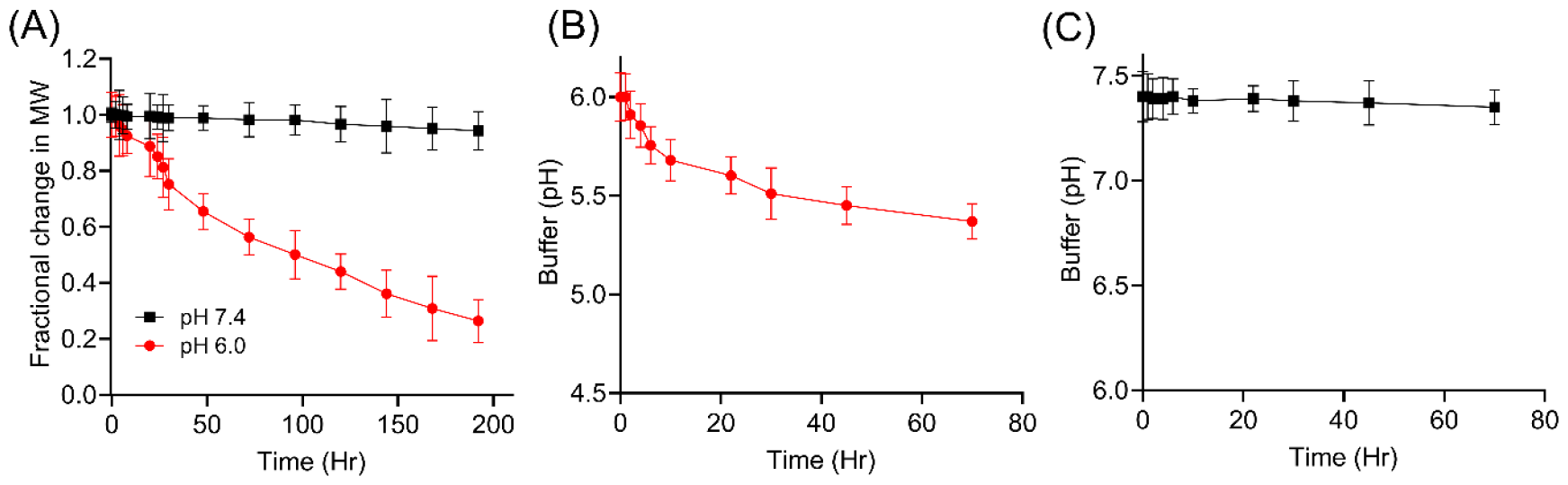
Degradation of AcNPs to impart acidifying property under mildly acidic condition. (A) Gel permeation chromatography analysis to determine the degradation rate of AcNPs under mildly acidic condition of pH 6.0 (red line) or neutral condition of pH 7.4 (black line). (B-C) Determination of the extent of buffer acidification by AcNPs under (B) mildly acidic condition of pH 6.0 (red line) and neutral condition of pH 7.4 (black line). Data are means ± SD of N=4 independent experiments.

**Fig. S2.**
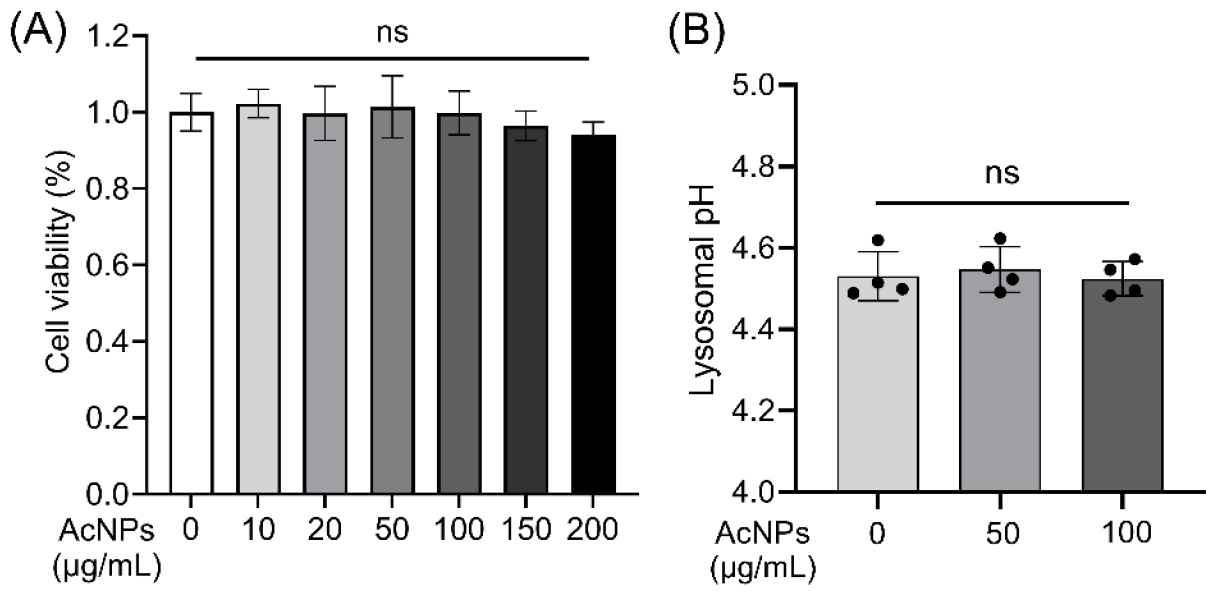
AcNPs are nontoxic and do not alter lysosomal pH of control SH-SY5Y cells. (A) Cytotoxicity assay illustrating that AcNPs at increasing concentration from 10 to 200 μg/mL are nontoxic to SH-SY5Y cells transfected with vector-only control. (B) AcNPs do not alter the basal level of lysosomal pH in SH-SY5Y cells transfected with vector-only control. Data are means ± SD of N=4 independent experiments and ns indicates non-significance using one-way ANOVA with post hoc Tukey’s test for multiple comparisons.

